# The Impact of Frailty Syndrome on Skeletal Muscle Histology: Preventive Effects of Exercise

**DOI:** 10.1101/2024.08.06.606836

**Authors:** Fujue Ji, Hae Sung Lee, Haesung Lee, Jong-Hee Kim

## Abstract

Aging-induced frailty syndrome significantly impairs skeletal muscle health, yet its impact on muscle histology remains unclear. This study investigates the histological alterations in muscle associated with frailty syndrome and evaluates the preventive effects of exercise. Mice were divided into groups based on age and condition, including an exercised group. Evaluated variables include body weight, lean mass ratio, myofiber size and number, extracellular matrix (ECM) content, and myosin heavy chain isoforms. Findings indicate that frailty syndrome increases body weight and ECM content, while reducing myofiber size and number, highlighting its negative impact on skeletal muscle histology. Notably, exercise effectively mitigated these adverse changes, suggesting its potential role in preventing skeletal muscle dysfunction associated with frailty syndrome.

## 1. Introduction

### Summary: Identification of Aging-Induced-Frailty Syndrome

Frailty syndrome is defined as a clinically recognizable state characterized by increased vulnerability resulting from age-associated decline in physiological reserve and function across multiple systems. This deterioration compromises an individual’s capacity to manage daily activities and respond to acute stressors effectively (Xue, 2011). Frailty syndrome encompasses three distinct states: pre-frailty, frailty, and frailty complications (Ahmed et al., 2007; Ji et al., 2024; Lang et al., 2009). Notably, individuals classified as pre-frail are at a heightened risk of progressing to frailty and subsequent complications, underscoring the downward spiral effect inherent in this syndrome (Ahmed et al., 2007; Gill et al., 2006).

### Summary: Why research relationship between frailty syndrome skeletal muscle

Skeletal muscle plays a critical role in whole-body metabolism, including energy metabolism, glucose regulation, and fat oxidation, making it an essential component of overall metabolic health (Franco-Romero et al., 2024; Mizushima, 2007). Impaired skeletal muscle health increases the risk of metabolic diseases such as diabetes and obesity, and even mortality (Argilés et al., 2016; Iizuka et al., 2014). Metabolic issues caused by the negative impacts of pre-frailty and frailty on skeletal muscle may also be one of the reasons contributing to the frailty complications. Although recent studies have identified multiple negative effects of frailty syndrome on skeletal muscle (Baumann et al., 2018, 2019; Ji et al., 2024; Kwak et al., 2020), the underlying specific details remain unclear. Studying histological changes may reveal the exact details of the negative effects of frailty syndrome on skeletal muscle.

### Summary: The correlation between histology and muscle healthy condition

Skeletal muscle health conditions are intricately linked to histological characteristics.. The mass and strength of skeletal muscle are directly indicated by the number and cross-sectional area (CSA) of myofibers (Verdijk et al., 2010). The metabolic function of skeletal muscle is determined by the distribution of myosin heavy chain (MHC) isoforms; MHC I fibers, which possess a higher number of mitochondria and blood vessels, primarily engage in aerobic metabolism (Schiaffino & Reggiani, 2011), while MHC II fibers contain more sarcoplasmic reticulum, providing greater instantaneous power (Schiaffino & Reggiani, 2011). The extracellular matrix (ECM) offers essential structural support for skeletal muscle, linking myofibers and playing a critical role in the transmission of muscular strength (Gillies & Lieber, 2011). Investigating these histological changes is essential for a comprehensive understanding of the mechanisms underlying skeletal muscle health conditions associated with frailty syndrome.

### Summary: Enhancing muscle histology through exercise

Endurance exercise has been extensively documented to enhance skeletal muscle health, primarily through its beneficial effects on muscle histology. Studies have shown that endurance exercise promotes skeletal muscle strength and power by increasing the number and CSA of myofibers (Bell et al., 2000; Green et al., 1999; Sanchez et al., 2005). Additionally, endurance exercise can remodel the ECM via regulated signaling pathways, specifically by modulating the matrix metalloproteinase family, which enhances the extracellular environment of myofibers and promotes overall muscle health (Das et al., 2008; Gibala et al., 2009). Furthermore, endurance exercise increases the proportion of MHC I isoforms, specifically by activating and modulating the AMP-activated protein kinase and CaN/NFAT Pathway, which enhances the mitochondrial biogenesis (Marcinko & Steinberg, 2014) and shift from MHC II isoforms to MHC I isoforms in response to sustained contractile activity (Yin et al., 2017). Those findings suggest that endurance exercise can effectively enhance skeletal muscle health conditions by improving its histological characteristics. However, the extent to which these histological improvements can prevent or mitigate the skeletal muscle health deterioration associated with frailty syndrome remains to be elucidated.

### Summary:Research purpose

Therefore, this study aims to investigate the histological characteristics of skeletal muscle in relation to frailty syndrome and to examine the preventive effects of endurance exercise on this condition. Understanding the histological changes in skeletal muscle may be crucial in diagnosing frailty syndrome-related diseases, thereby aiding physicians in identifying and assessing pathological changes. Furthermore, the beneficial effects of endurance exercise on preventing frailty-related diseases provide scientific support for the development of future exercise prescriptions.

## 2. Methods

### 2.1 Animals and experimental design

Male C57BL/6N mice were used as experimental subjects and allocated into five groups: young (2 months old, YM, n=10), adult (13 months old, AM, n=10), old (22 months old, OM, n=33), frailty syndrome (22 months old, FS, n=13), and endurance exercise (22 months old, EOM, n=10) (Figure 1). The room temperature (22 ± 2 °C), humidity (50–60%), and light (12/12 h light/dark cycle) were controlled. The mice had ad libitum access to food, water, and activity. All procedures were approved by the Institutional Animal Care and Use Committee (IACUC) of Hanyang University (HYU 2017-0265A; 2019-0017A) and adhered to relevant guidelines and regulations, in compliance with ARRIVE guidelines (https://arriveguidelines.org).

**Figure 1.**
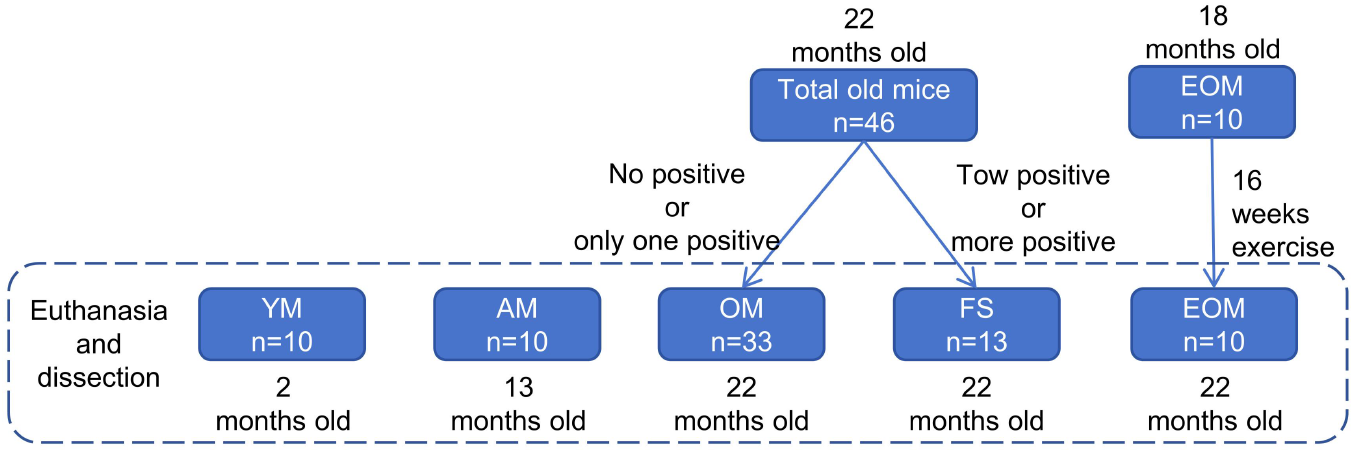
Animals and experimental design.

### 2.2 Screening of frailty syndrome mouse model

The screening criteria for the frailty syndrome mouse model employed in this study have been described in detail in previous research (Baumann et al., 2018, 2019; Ji et al., 2024; Kwak et al., 2020). Briefly, 22 months old mice that fell within the bottom 20% for measures of slowness (rota-rod test), weakness (grip strength test), poor endurance (treadmill test), or low physical activity (voluntary wheel test), or those in the top 20% for body weight within our cohort, were identified as positive markers for frailty syndrome. These criteria were established to determine the cut-off values for frailty syndrome (Table 1).

**Table 1.**
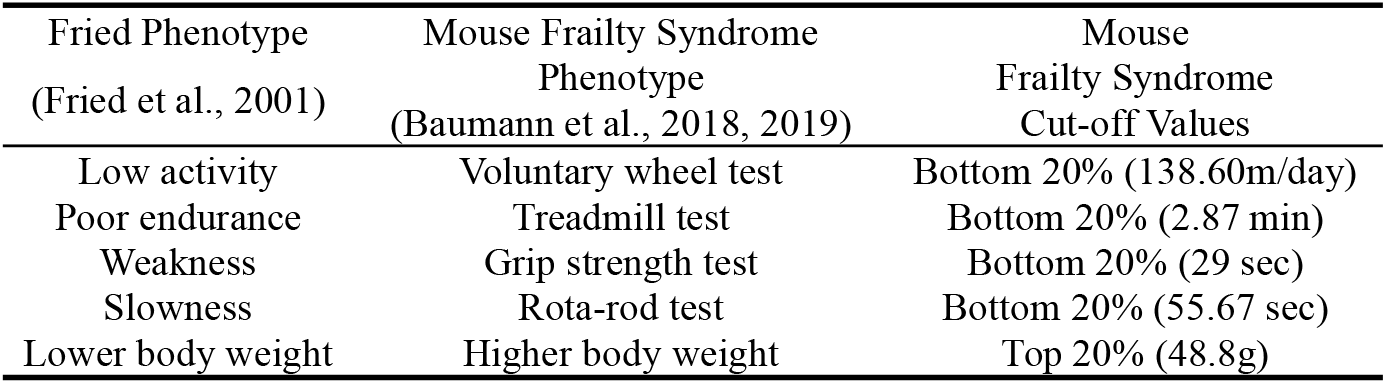
Frailty syndrome criteria and cut-off values.

Mice with two or more positive markers were classified as exhibiting frailty syndrome, and those with one or no positive marker was categorized as non-frailty syndrome (Baumann et al., 2018, 2019; Ji et al., 2024; Kwak et al., 2020) (Figure 2).

**Figure 2.**
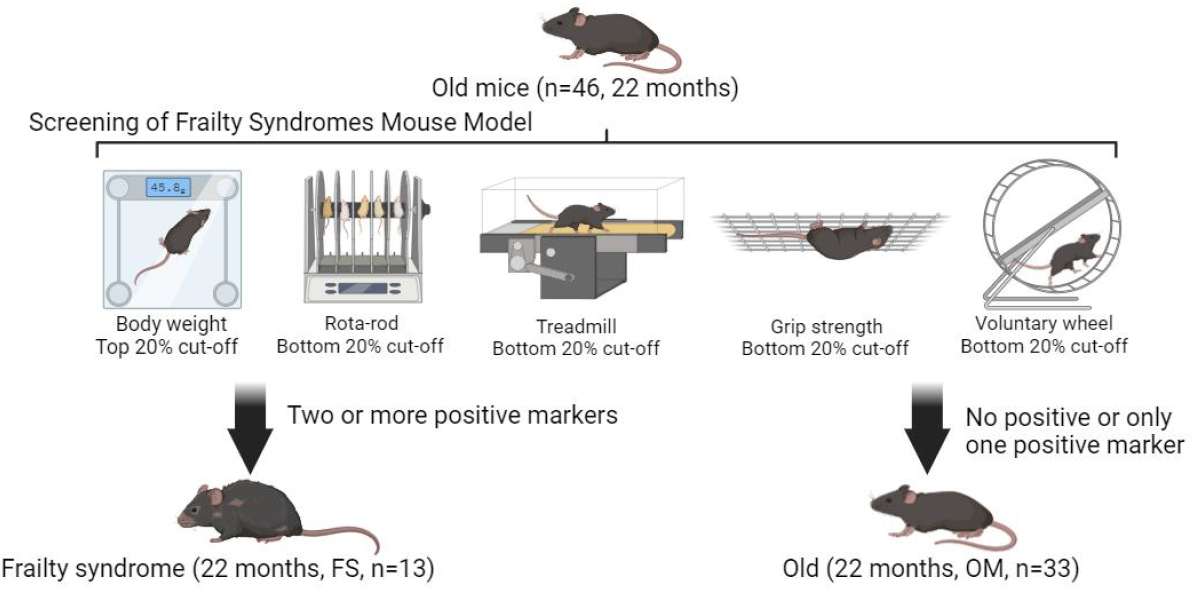
The frailty syndrome mouse model identification process.

#### Body weight

Body and skeletal muscle raw weights were obtained on an electronic scale (CS200; OHAUS, Parsippany, NJ).

#### Walking speed

Walking speed was assessed using the Rota-Rod test (Ji et al., 2024). A three-day adaptation phase included a pre-test exercise at 5 rpm for 1 minute, conducted once daily. The formal test employed an acceleration mode, progressively increasing speed from 5 to 50 rpm over a span of 5 minutes (Model 76-0770, Harvard Apparatus Inc., Holliston, MA, USA). The latency to fall from the apparatus was measured. Each mouse underwent three trials with a ten-minute interval between tests, and the best performance among the three trials was used as the final outcome measure.

#### Grip strength

Grip strength was evaluated using the inverted-cling grip test (Ji et al., 2024). Mice underwent an adaptation phase with a daily trial for three days before the official test. For the official test, each mouse was placed at the center of a wire mesh screen, and a timer was started. The screen was then inverted over the course of 2 seconds, positioning the mouse’s head downward, and held 40-50 cm above a padded surface. The time until the mouse released its grip and fell was recorded. This measurement was repeated three times with ten-minute intervals between tests, and the longest duration was taken as the final measurement.

#### Endurance

Endurance capacity was assessed using a treadmill test (Ji et al., 2024). Mice underwent an adaptation period for three days prior to the official test, running at a speed of 5 cm/s on a 0-degree incline for 5 minutes daily. During the official test, the treadmill speed began at 5 cm/s and increase by 1 cm/s every 20 seconds, maintaining a 0-degree incline. The test concluded when the mouse contacted the shock pad (set at 0.5 mA) three times.

#### Physical activity

Physical activity was measured using the voluntary wheel test (Ji et al., 2024). Running distance was monitored using a voluntary wheel apparatus (MAN86130, Lafayette Instrument Company, Lafayette, IN, USA), with each wheel rotation equating to a distance of 0.4 meters. The average running distance over a 5-day period was recorded for each mouse.

### 2.3 Exercise protocol

The EOM group (18 months old) underwent 16 weeks of endurance exercise. At the end of the endurance exercise period (22 months old), they were euthanized. The EOM group was subjected to endurance exercise on a treadmill, with a preparatory adaptation phase conducted one week prior to the main experiment, during which all aged mice underwent three sessions of acclimatization at an intensity of 5 m/min for 20 minutes per session. Following the start of the formal experiment, mice exercised for 60 minutes, three times a week, for 16 weeks. Exercise intensity was gradually increased from 4 to 16 m/min per week and subsequently maintained, with the treadmill incline set at 0°. Instead of using electrical stimulation, the mice were gently guided to run naturally by using a wooden stick as needed (Figure 3).

**Figure 3.**
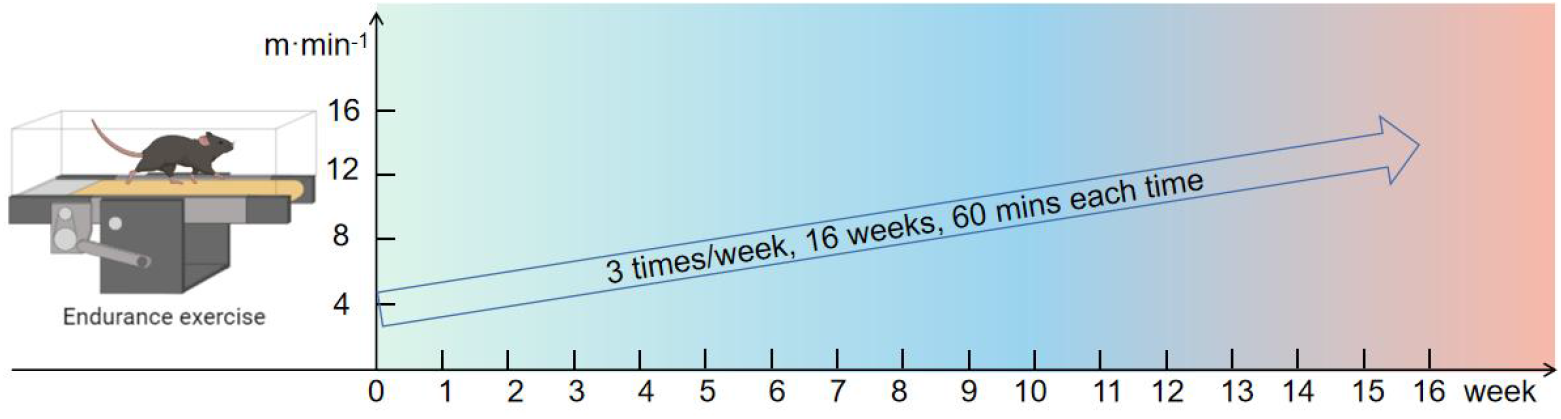
Endurance exercise protocol for EOM

### 2.4 Preservation and preparation of skeletal muscle

After the mice were euthanized, the tibialis anterior (TA), quadriceps (QD), and plantaris (PL) were carefully removed. The left leg muscles were fixed in 10% formalin for 24 hours, then dehydrated, and embedded in paraffin. The sections were prepared for future hematoxylin and eosin (H&E) and Masson’s trichrome staining. The right leg muscles were weighed, frozen in liquid nitrogen, and subsequently stored at –80 °C for subsequent MHC isoform analyses.

### 2.5 Masson’s trichrome staining and analysis

Paraffin-embedded tissues were sectioned at 5 μm. After deparaffinization and rehydration, the tissue sections were stained using Masson’s trichrome staining kit (G1340, Beijing Solarbio Science & Technology Co., Ltd.). The sections were subsequently placed in xylene and alcohol for dehydration and transparency, and finally sealed with neutral resin. Images were acquired using a slide scanner (Axio Scan. Z1, Zeiss) at 200x magnification. For each mouse, a single section of each skeletal muscle was randomly photographed to obtain 5 images, each covering an area of 0.271 square millimeters. The average of these five images was used to represent the ECM value of that skeletal muscle for the mouse. ImageJ software was used for analysis. A threshold was established to remove the blank background, the ECM signal was isolated using color deconvolution and thresholding, and the area of the ECM signal was subsequently compared with the net area of the muscle section using ImageJ software.

### 2.6 H&E staining and analysis

H&E staining was performed on paraffin-embedded tissue sections after deparaffinization and rehydration, as previously described by Robert D. Cardiff (Almond & Enser, 1984), with slight modifications. Briefly, rehydrated tissue sections were stained using ClearView™ hematoxylin (MA0101010, StatLab) for 3 minutes, rinsed in tap water for 30 seconds, differentiated in 1% hydrochloric acid alcohol for 5 seconds, washed in tap water for 3 minutes, and stained in ClearView™ eosin (MA0101015, StatLab) for 1 minute. They were subsequently dehydrated, rendered transparent, and sealed with neutral resin. Images were acquired using a slide scanner (Axio Scan. Z1, Zeiss) at 200x magnification. For each mouse, a single section of each skeletal muscle was randomly photographed to obtain 5 images, each covering an area of 0.271 square millimeters. The average of these five images was used to represent the CSA and fiber number of that skeletal muscle for the mouse. The CSA and fiber number were automatically analyzed using Cellpose (Stringer et al., 2021; Waisman et al., 2021).

### 2.7 Electrophoretic separation of MHC isoforms

The MHC isoform composition of each fiber was determined using SDS–PAGE and silver staining. Skeletal muscle was extracted for 60 min on ice in 50 vol buffer (0.1 M sodium phosphate, pH 7.3), to which a 1:100 protease + phosphatase inhibitor cocktail (WSE-7420, ATTO) was added. The total protein concentration was subsequently determined using the Pierce™ BCA Protein Assay Kit (23227, Thermo). Samples were diluted in 5× sample buffer (25% β-MEtOH [β-ME], 11.5% SDS, 50% glycerol, 312.5 mM Tris, pH 6.8, and 0.05% bromophenol) and dH_2_O to achieve a final protein concentration of 0.266 μg/μL. The samples were subsequently boiled for 10 min, and 6 μg (30 μL) of protein was loaded onto a mini gel system (Mini Protean III: 8.3 × 7.3 cm). MHC isoforms were separated using 8% SDS–PAGE with 30% glycerol (Talmadge & Roy, 1993). The lower running buffer comprised 0.05 M Tris (base), 75 mM glycine, and 0.05% w/v SDS. The upper running buffer was 6× the concentration of the lower running buffer, and β-ME was added (final concentration: 0.2% v/v) (Kohn & Myburgh, 2006). After sample loading, electrophoresis was performed at a constant voltage of 140 V for 28 h. The temperature of the buffer was maintained at 4 °C during electrophoresis. After electrophoresis, the gels were stained with a silver staining kit (24612, Thermo), and the bands were quantified via densitometry using Image J software.

### 2.8 Statistical analysis

Analyses were performed using GraphPad Prism (version 9) software. One-way analysis of variance and Tukey’s post-hoc test were used to compare the mean difference among groups. When the conditions of normality or homogeneity were not met in comparisons of three or more groups, the Kruskal–Wallis test along with Dunn’s post-hoc comparison was employed. All results are expressed as the mean ± standard error of the mean (SEM). Statistical significance was set at p < 0.05. Asterisks indicate the following: *p < 0.05, **p < 0.01, ***p < 0.001, and ****p < 0.0001.

## 3. Results

### 3.1 Endurance exercise prevents frailty syndrome by reducing body weight

Aging leads to an increase in body weight, and frailty syndrome further exacerbates this phenomenon (P < 0.05) (Figure 4A). Aging also reduces the lean mass ratio (P < 0.05), but frailty syndrome does not (P > 0.05) (Figure 4B-D). After 16 weeks of endurance exercise, there was a significant reduction in body weight, compared to the OM and FS groups (P < 0.05) (Figure 4A). The lean mass ratio also significantly increased (P < 0.05) (Figure 4B-D). These data suggest that aging leads to weight gain, and frailty syndrome exacerbates this phenomenon but does not significantly alter the lean mass ratio. Importantly, 16 weeks of endurance exercise can prevent frailty syndrome by reducing body weight.

**Figure 4.**
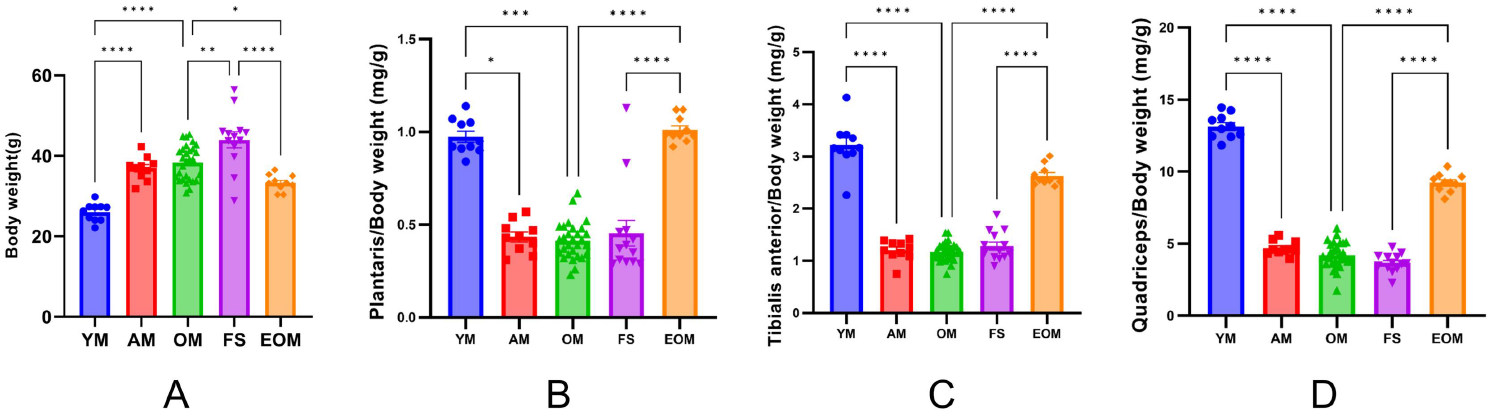
Body weight and lean muscle ratio. (A) Body weights of the different groups of mice. (B, C, and D) Plantaris, tibialis anterior, and quadriceps to body weight ratios. YM = 2 months old, n = 10; AM = 13 months old, n = 10; OM = 22 months old, n = 33; FS= 22 months old, n = 13; and EOM = 22 months old, n = 10. All data are presented as the mean ± SEM (*p<0.05, **p<0.01, ***p<0.001, ****p<0.0001).

### 3.2 Endurance exercise prevents frailty syndrome by reducing ECM content

Experimental results show that the ECM content of the PL and QD increases with age (P < 0.05), while the TA does not appear to be affected by age (P > 0.05) (Figure 5B-D). The impact of frailty syndrome on skeletal muscle ECM content is significant. In the PL, TA, and QD, the ECM content in the FS group is significantly larger than in the OM group (P < 0.05) (Figure 5B-D). Additionally, after 16 weeks of endurance exercise, the ECM in the EOM group is significantly lower than in the OM and FS groups (P < 0.05) (Figure 5A-D). These findings indicate that aging is a key factor leading to abnormal ECM content accumulation in skeletal muscles, and this negative effect is significantly exacerbated by frailty syndrome. Importantly, 16 weeks of endurance exercise can prevent frailty syndrome by inhibiting abnormal ECM content accumulation.

**Figure 5.**
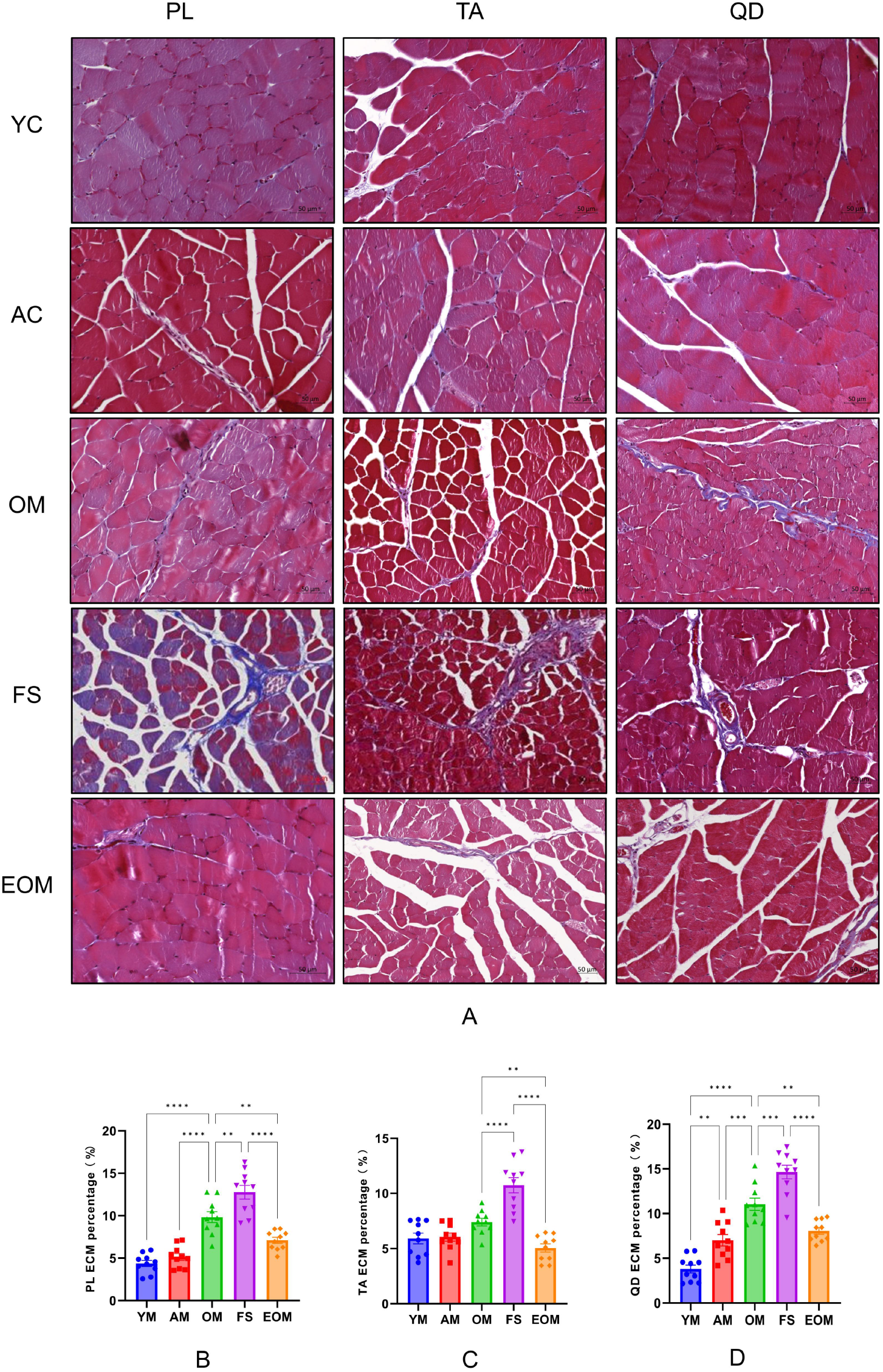
Representative images of Masson’s trichrome staining of the plantaris, tibialis anterior, and quadriceps extracellular matrixes of male C57BL/6N mice at different stages. (A) Representative images of Masson’s trichrome staining. (B) Changes in the extracellular matrix of the plantaris in male C57BL/6N mice. (C) Changes in the extracellular matrix of the tibialis anterior in male C57BL/6N mice. (D) Changes in the extracellular matrix of the quadriceps in male C57BL/6N mice. YM = 2 months old, n = 10; AM = 13 months old, n = 10; OM = 22 months old, n = 10; FS= 22 months old, n = 10; and EOM = 22 months old, n = 10. The extracellular matrix content is indicated in blue and the myofibers in red. All data are presented as the mean ± SEM (*p<0.05, **p<0.01, ***p<0.001, ****p<0.0001). Scale bar=50 μm.

### 3.3 Endurance exercise prevents decreased CSA due to frailty syndrome

The effects of aging on the CSA of different skeletal muscles vary, which may be determined by the functional roles in daily activities. We observed that in the TA and QD, the CSA of the FS group was significantly lower than that of the OM group (P < 0.05) (Figure 6C and D). In contrast, in the PL, the CSA of the FS group was significantly higher than that of the OM group (P < 0.05) (Figure 6B). Additionally, after 16 weeks of endurance exercise, the CSA of the EOM group was significantly higher than that of the OM and FS groups in TA and QD, but not PL (P < 0.05) (Figure 6B-D). These data demonstrate that the impact of frailty syndrome on skeletal muscle CSA varies; 16 weeks of endurance exercise can partially prevent the decline in skeletal muscle CSA caused by frailty syndrome.

**Figure 6.**
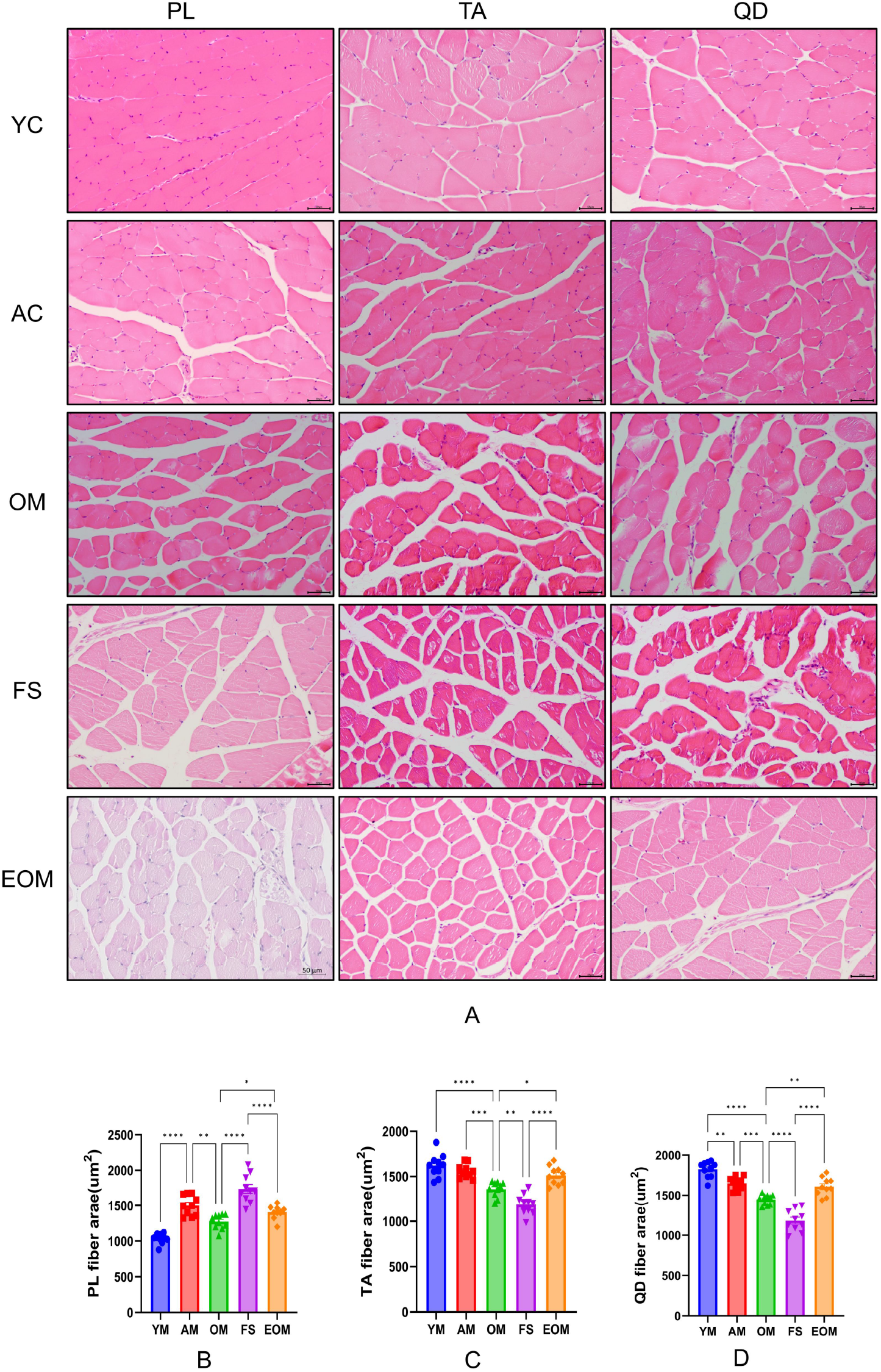
Representative images showing H&E staining of plantaris, tibialis anterior, and quadriceps cross-sectional areas in male C57BL/6N mice at various stages. (A) Representative images of H&E staining. (B) Changes in the cross-sectional area of the plantaris in male C57BL/6N mice. (C) Changes in the cross-sectional area of the tibialis anterior in male C57BL/6N mice. (D) Changes in the cross-sectional area of the quadriceps in male C57BL/6N mice. YM = 2 months old, n = 10; AM = 13 months old, n = 10; OM = 22 months old, n = 10; FS= 22 months old, n = 10; and EOM = 22 months old, n = 10. All data are presented as the mean ± SEM (*p<0.05, **p<0.01, ***p<0.001, ****p<0.0001). Scale bar=50 μm.

### 3.4 Endurance exercise prevents decreased myofiber number due to frailty syndrome

Data show that aging leads to a significant reduction in the myofiber number and frailty syndrome further exacerbates this decrease (P < 0.05) (Figure 7A-C). Additionally, 16 weeks of endurance exercise EOM increased the myofiber number, significantly surpassing both the FS and OM groups (P < 0.05) (Figure 7A-C). These experimental results reveal that frailty syndrome reduced myofiber number; 16 weeks of endurance exercise can prevent the negative effects of frailty syndrome on skeletal muscle histology by increasing the number of myofiber.

**Figure 7.**
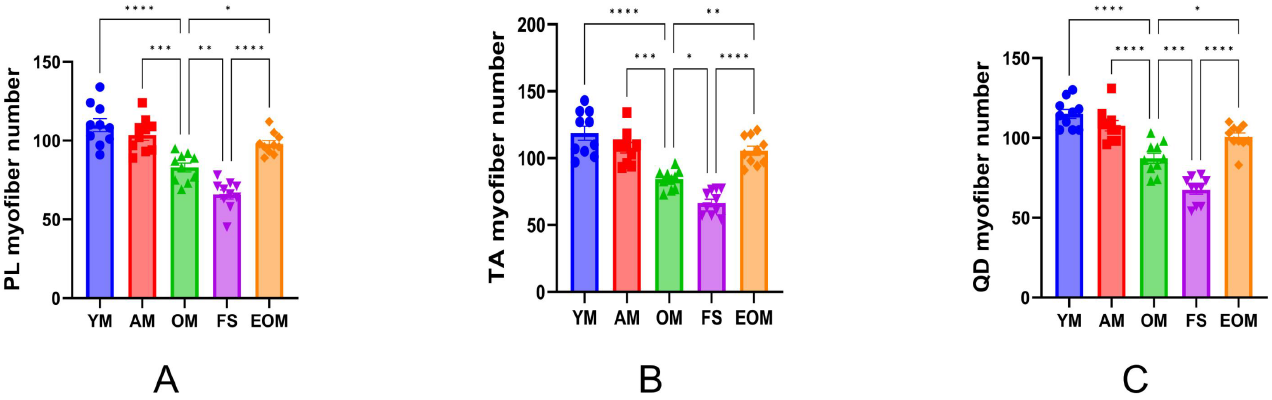
Changes in the myofiber number of the plantaris, tibialis anterior, and quadriceps muscles in male C57BL/6N mice at different stages. (A) Changes in the myofiber number of the plantaris in male C57BL/6N mice. (B) Changes in the myofiber number of the tibialis anterior in male C57BL/6N mice. (C) Changes in the myofiber number of the quadriceps in male C57BL/6N mice. YM = 2 months old, n = 10; AM = 13 months old, n = 10; OM = 22 months old, n = 10; FS= 22 months old, n = 10; and EOM = 22 months old, n = 10. All data are presented as the mean ± SEM (*p<0.05, **p<0.01, ***p<0.001, ****p<0.0001).

### 3.5 Frailty syndrome does not affect MHC isoforms

We used an anti-MHC (Myosin 4 Monoclonal Antibody, Thermo, #14-6503-82) to examine the band positions in the gel, confirming that the bands observed in silver staining correspond to MHC (S1). Aging leads to an increase in the proportion of MHC I and a decrease in the proportion of MHC IIX in TA (P < 0.05) (Figure 8). Similarly, in QD, aging significantly increases the proportion of MHC I (P < 0.05) (Figure 8). We also found that frailty syndrome does not significantly affect the proportion of MHC in skeletal muscle (P > 0.05) (Figure 8). 16 weeks of endurance exercise resulted in a significant decrease in the proportion of MHC IIB only in TA (P < 0.05) (Figure 8). These data indicate that frailty syndrome does not affect the proportion of MHC in skeletal muscle.

**Figure 8.**
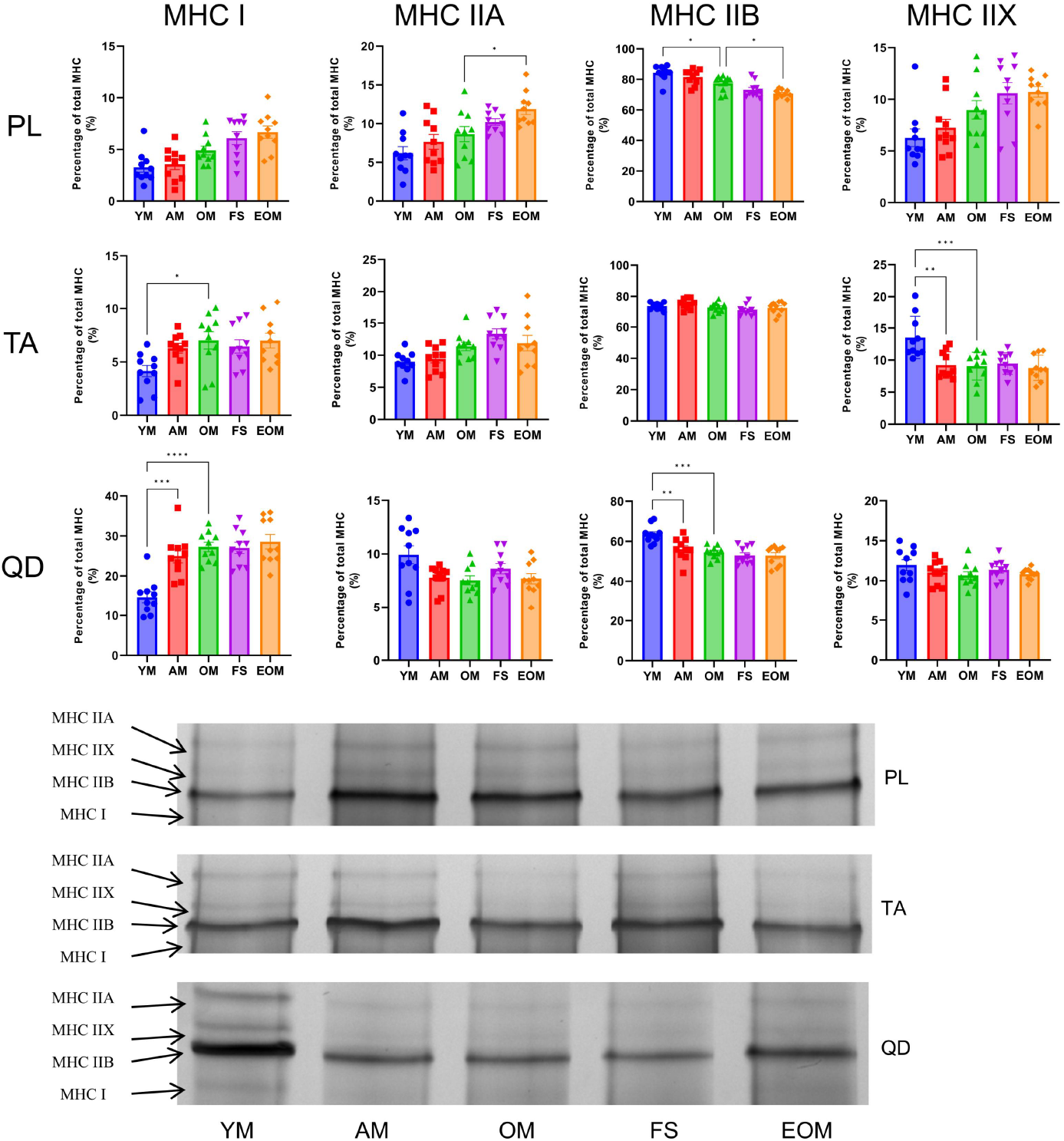
Changes in the myosin-heavy-chain isoform percentage of the plantaris, tibialis anterior, and quadriceps in male C57BL/6N mice at different stages. YM = 2 months old, n = 10; AM = 13 months old, n = 10; OM = 22 months old, n = 10; FS= 22 months old, n = 10; and EOM = 22 months old, n = 10. All data are presented as the mean ± SEM (*p<0.05, **p<0.01, ***p<0.001, ****p<0.0001).

## 4. Discussion

Although the negative effects of frailty syndrome on skeletal muscle health have been widely demonstrated (Baumann et al., 2018, 2019; Ji et al., 2024; Kwak et al., 2020), the specific details and potential preventive measures remain unclear. Therefore, there is an urgent need to uncover the details of the negative effects of frailty syndrome on skeletal muscle and identify preventive measures, which could provide valuable insights for the treatment and prevention of aging-related diseases. We found that frailty syndrome leads to increasing body weight and ECM accumulation, while decreasing CSA and myofiber number. Importantly, endurance exercise can prevent the negative effects of frailty syndrome on skeletal muscle by improving histological health condition. These findings provide scientific data for the prevention, diagnosis, treatment, and post-recovery evaluation of frailty syndrome in the future.

### 4.1 Appropriateness of a frailty syndrome mouse model

In our frailty syndrome mouse model, we followed the selection method described by Thompson et al. (Baumann et al., 2018, 2019; Kwak et al., 2020). In animal studies on frailty syndrome, this modeling method is widely used (Baumann et al., 2018, 2019; Ji et al., 2024; Kwak et al., 2020; Marcozzi et al., 2023; Yilmaz et al., 2024). Differing from the Fried frailty syndrome phenotype, instead of unintentional weight loss, we determined that old mice with a high body weight, ranking in the top 20%, is a positive marker for frailty syndrome. Briefly, there are several reasons. Firstly, high body weight has a significant negative impact on the lifespan of mammals (Economos, 1980). A typical example is the Ames dwarf, which has a longer lifespan (Bartke et al., 2003; Brown-Borg et al., 1996), and calorie restriction, which leads to lower body weight and increases both average and maximum lifespan (Masoro, 1993; Weindruch, 1989). Additionally, an increase in body weight is positively correlated with mortality in mice (Miller et al., 2000; Miller et al., 2002). High body weight poses a greater threat to mice than low body weight, which aligns more closely with the negative health impacts characteristic of frailty syndrome. For a detailed explanation, refer to the excellent paper by Thompson et al. (Baumann et al., 2020). The 20% cut-off point was set based on the percentiles used in the diagnosis of frailty syndrome in human study (Fried et al., 2001). The 20% cut-off point is widely recognized in research for diagnosing frailty syndrome in mice (Baumann et al., 2018, 2019; Ji et al., 2024; Kwak et al., 2020). The specific selection process for the frailty syndrome mice is detailed in the supplementary materials (S1).

Selecting 22 months old mice as experimental subjects is intended to better simulate the characteristics of human frailty syndrome. 22 months old mice roughly corresponds to a human aged 60-65 years (Dutta & Sengupta, 2016; Justice et al., 2016), which is the period when frailty syndrome typically emerges during normal aging (Calado et al., 2016; Fried et al., 2001). Moreover, because frailty syndrome is thought to be reversible, this age provides adequate time to implement possible life-changing interventions, such as endurance exercise.

We want to emphasize that the onset of frailty syndrome is a continuous process (Ahmed et al., 2007), currently broadly categorized into three stages: pre-frailty, frailty, and frailty complication (Ahmed et al., 2007; Lang et al., 2009). Research has shown that patients in the pre-frailty stage are more likely to progress to the frailty and frailty complication stages compared to non-pre-frailty patients (Gill et al., 2006). Additionally, there is potential for reversal during the pre-frailty stage (Gill et al., 2006). The final stage of frailty syndrome involves complications that ultimately lead to patient mortality (Robertson & Montagnini, 2004). To our knowledge, this is the first study investigating the impact of frailty syndrome on skeletal muscle histology. This research will provide scientific data to support future exploration of the frailty syndrome in skeletal muscle.

### 4.2 Lean mass ratio in frailty syndrome

Although the lean mass ratio decreased with age, no significant difference was noted between the OM and FS groups. This may be related to the high body weight observed in the frailty syndrome mouse model. High body weight potentially causes ectopic fat deposition in skeletal muscle, known as myosteatosis (fat infiltration of skeletal muscle); and increased ECM (see below) in skeletal muscle also potentially exacerbates the risk of myosteatosis. Therefore, we believe that the fat infiltration caused by high body weight increases the skeletal muscle mass and this is the main reason why the lean mass ratio does not decrease in frailty syndrome. Moreover, the skeletal muscle of mice with frailty syndrome possibly also undergo prolonged heavy weight-bearing to maintain basic vital activities (e.g., postural maintenance). The compensatory effects of weight gain-induced increases in skeletal muscle mass and strength have also been identified in previous studies (Abdelmoula et al., 2012; Duché et al., 2002). Notably, endurance exercise has been shown to beneficial effect in preventing frailty syndrome by significantly reducing body weight and increasing the lean mass ratio.

### 4.3 ECM in frailty syndrome

The ECM of the FS group was significantly increased than that of OM, which to our knowledge is the first time it has been reported. While it is not clear how frailty syndrome affects ECM, some studies have found that both aging (Clemens et al., 2021; Liou et al., 2013) and high body weight (Pincu et al., 2015) lead to increased ECM. The abnormal increase in the ECM in frailty syndrome might have resulted from a combination of aging and high body weight. Specifically, aging and high body weight lead to the excessive accumulation of ECM components such as collagen and fibrinogen through oxidative stress and inflammatory responses (Ruiz-Ojeda et al., 2019). These changes are mediated by alterations in the activity of matrix metalloproteinases and the activation of hypoxia-inducible factors (Brown et al., 2016). Moreover, excessive ECM accumulation accelerates fat infiltration into skeletal muscle, which may be responsible for the abnormal lean mass ratio in frailty syndrome. The abnormal increase in the ECM may be one of the skeletal muscle histological characteristics of frailty syndrome.

Endurance exercise can prevent frailty syndrome by reducing the skeletal muscle ECM in aging individuals. Numerous reports state that endurance exercise can decrease ECM content and improve its health status (Csapo et al., 2020; Song et al., 2006). A healthy ECM content not only efficiently transmits skeletal muscle strength (Csapo et al., 2020) and provides mechanical support to nerves and blood vessels (Sheard & Anderson, 2012) but also promotes the proliferation and differentiation of satellite cells (Csapo et al., 2020). An unhealthy ECM state may the one of the negative factors contributing to or causing frailty syndrome. Based on our data, endurance exercise may be one of the methods of preventing frailty syndrome by remodeling and reducing the ECM content.

### 4.4 CSA in frailty syndrome

Unexpectedly, frailty syndrome mice exhibited an abnormal increase in CSA of PL. To the best of our knowledge, this is the first time it has been found in the PL in frailty syndrome. We hypothesis that the loss and reduced proportion of smaller myofiber in the PL lead to an increase in the mean area or a compensatory mechanism that attempts to compensate for the loss of skeletal muscle strength to maintain the basal vital activity of the organism (Sayed et al., 2016; Sheard & Anderson, 2012). Moreover, during the experiment, we observed that mice with frailty syndrome had increased resting time and decreased physical activity, exacerbating their sedentary status (S1). The PL is more involved in postural maintenance than the QD and TA (Akay et al., 2014; Charles et al., 2018), possibly explaining the abnormal increase in the CSA of the PL in frailty syndrome because of compensatory effects, a phenomenon not apparent in the QD and TA.

The significant reduction in the CSA of QD and TA may result from multiple negative factors induced by frailty syndrome. Frailty syndrome leads to a decrease in myofiber CSA through various molecular biological mechanisms. Firstly, autophagy dysregulation associated with frailty syndrome affects the clearance of skeletal muscle proteins, leading to a decline in myofiber CSA (Aas et al., 2019; Mammucari et al., 2007). Additionally, the imbalance between protein synthesis and degradation due to aging induced-frailty syndrome, results in a significant decline in myofiber CSA (Grimby, 1995). Finally, increased levels of oxidative stress and chronic inflammation in frailty syndrome elderly individuals damage skeletal muscle cells and tissues, further contributing to the decline in CSA (Lang et al., 2010). Therefore, we believe that the significantly reduced CSA is one of the skeletal muscle histological characteristics of frailty syndrome.

Our experimental results indicate that endurance exercise significantly increases myofiber CSA. This finding aligns with previous studies (Clark et al., 1989; Coggan et al., 1992; Lewis et al., 2015; Wang et al., 2023). Endurance exercise promotes the increase in myofiber CSA through various molecular biological mechanisms. Even endurance exercise, which is traditionally not considered to cause significant muscle hypertrophy, also can lead to increases in myofiber CSA when performed over a long period and at appropriate intensities (Konopka & Harber, 2014). Specifically, endurance exercise can activate the AKT and mTOR signaling pathways, promoting skeletal muscle hypertrophy, particularly during the post-exercise recovery period (Camera et al., 2010). Additionally, endurance exercise increases the expression of vascular endothelial growth factor, promoting capillary formation in skeletal muscle, enhancing oxygenation, and supporting the growth and metabolic needs of myofiber (Kon et al., 2014). These mechanisms collectively contribute to the significant increase in myofiber CSA through endurance exercise. Therefore, we believe that the CSA reduction caused by frailty syndrome can be prevented through endurance exercise.

### 4.5 Myofiber Number in Frailty Syndrome

Our results indicated that myofiber number was significantly reduced in FS compared with that in the OM group, possibly because of the simultaneous activation of apoptotic pathways by high body weight (Heo et al., 2018) and aging (Ziaaldini et al., 2015). Specifically, aging and high body weight lead to an increase in pro-apoptotic factors (e.g., P53 and Bax) and a decrease in anti-apoptotic factors (e.g., Bcl-2) in skeletal muscle, potentially inducing a decrease in myofiber number with aging and high body weight (Agrawal et al., 2024; Lin et al., 2017; Yahagi et al., 2003; Ziaaldini et al., 2015). Therefore, we consider the reduction in myfiber number to be one of the skeletal muscle histological characteristics of frailty syndrome.

After 16 weeks of endurance exercise, the myofiber number significantly increased, consistent with the findings of previous studies (Gonyea et al., 1986; Hoffman et al., 2013; Luo et al., 2023; Waters et al., 2004). Endurance exercise may promote the increase in myofiber number through various molecular biological mechanisms. Firstly, as described above, endurance exercise increases the rate of skeletal muscle protein synthesis, which is one of the reasons for the rise in myofiber number. Secondly, calmodulin-dependent kinases, can decode frequency-dependent information to activate adaptive responses, thus promoting the increase in myofiber number (Chin, 2005). Finally, autophagy is critical in supporting skeletal muscle adaptive changes. Endurance exercise increases autophagic flux, helping to clear damaged organelles and proteins (Sanchez et al., 2014), thereby providing space and raw materials for the increase in myofiber number. These mechanisms collectively contribute to a significant increase in the myofiber number. Therefore, the reduction in myofiber number caused by frailty syndrome can be prevented through endurance exercise.

### 4.6 MHC isoforms in frailty syndrome

MHC isoforms potentially reflect skeletal muscle physiological and functional characteristics. We found that aging leads to reduce the percentage of fast MHC isoforms (e.g., IIb and IIx), exhibiting consistency with mainstream findings (Balagopal et al., 2001; Lopez et al., 2000; Uchitomi et al., 2019). Recent studies on MHC isoforms in mice with frailty syndrome are inadequate, and our data did not reveal significant differences in it between OM and SF group. Aging leads to decreased fast MHC isoforms and related-gene expression (Balagopal et al., 2001; Ciciliot et al., 2013; Uchitomi et al., 2019). Despite some controversy, studies have found that high body weight increases fast MHC isoforms and related-gene expression in humans and rodents (Almond & Enser, 1984; Houmard et al., 2002; Lillioja et al., 1987). Therefore, we hypothesis that the unchanged MHC isoforms percentage in frailty syndrome emanates from the combined effects of aging and high body weight.

Only the slow MHC isoforms percentage of the PL increased significantly after 16 weeks of endurance exercise. Fiber diameter and area were negatively correlated with oxidative capacity, with small myofibers possessing higher oxidase activity and mitochondrial density than large, regardless of fiber type (Umek et al., 2019; van Wessel et al., 2010). Based on our data, the mean CSA of the PL was smaller than that of the QD and TA, potentially explaining the changes in the MHC isoforms percentage of the PL after endurance exercise. This is because more oxidase activity is activated in the PL after endurance exercise than in the QD and TA, thus increasing oxidative fibers and slowing MHC isoforms percentage.

## 5. Limitations

In the histological experiments, skeletal muscle was not examined at high resolution using transmission electron microscopy (TEM). TEM allows for the examination of ultrasmall changes in skeletal muscle fibers and organelles in frailty syndrome mice. The present study investigated the histological characteristics of frailty syndrome mice. In the future, the specificity of protein and gene expression in mice with frailty syndrome should be further investigated.

## 6. Ethical approval and consent to participate

All the procedures followed in this experiment were approved by the Institutional Animal Care and Use Committee (IACUC) of Hanyang University (HYU 2017-0265A; 2019-0017A).

## 7. Consent for publication

Not applicable.

## 8. Availability of data and materials

The data used to support the findings of this study are presented here. Any further data requirements are available from the corresponding author upon request.

## 11. Author’s contributions

*Fujue Ji:* Conceptualization, data curation, formal analysis, investigation, methodology, writing—original draft.

*Hae Sung Lee:* Data curation and investigation.

*Haesung Lee:* Formal analysis, and methodology.

*Jong-Hee Kim:* Conceptualization, supervision, funding acquisition, writing — review and editing.

## 12. Acknowledgements

We thank Ran Tao (Beijing Solarbio Science & Technology Co., Ltd., China) for experiment-related technical assistance.

We thank the support of Skill Learning from Kaixin Doctor and MASCU (Medical Association with Science, Creativity, and Unity), Inc., Shenzhen, China.

